# Molecular Insights into Solvent-Mediated Stabilization and Aggregate Suppression during Refolding of Recombinant Leucyl Aminopeptidase

**DOI:** 10.64898/2026.06.12.732004

**Authors:** Diptesh Das, Jai Kumar Kaushik

## Abstract

Production of recombinant proteins frequently yields inclusion bodies that must undergo refolding to yield active protein. Here, we optimized the refolding conditions for the recombinant leucyl aminopeptidase (rPepL) from Lactocaseibacillus casei expressed in inclusion bodies from E. coli. Several chemical additives were assessed for how well they facilitated an increase in refolding efficiency. The best, 0.5 M L-arginine, yielded 50.8% refolding. The addition of stabilizers, such as sucrose and glycerol, with L-arginine further increased yields to 85%. Urea at lower concentrations (0.25-0.5 M) also facilitated an increase in the refolding yield when co-added with L-arginine, whereas guanidinium chloride inhibited it. Sugars and polyols exhibited dose-dependent effects, with ranges for optima also defined. Fluorescence spectroscopy verified enhancements in the refolding under the optimized conditions. Molecular dynamics simulation under mixed solvent conditions provided atomic insights about stabilizing interactions that are likely to facilitate increased refolding. The results show that a series of aggregation suppressors and protein stabilizers can, in a collaborative way, increase the refolding efficiency for the recombinant proteins from the inclusion bodies. The protocol with the optimization using the additives L-arginine, sucrose, and glycerol is an efficient method for the production of active rPepL.

This article outlines the best refolding method to recover recombinant leucyl aminopeptidase from inclusion bodies of E. coli using L-arginine combined with sucrose and glycerol. The combined experimental observations and computational simulations elucidate the molecular process of additive-induced stabilization, which elucidates how aggregation inhibition and hydrogen-bonded stabilization act synergistically. The results presented herein answer both mechanistic understanding and experimental guidance for improving protein refolding.

## 1. Introduction

Availability of large amounts of target protein is a major obstacle in biopharmaceuticals manufacturing, therapeutic drug design, structural biology. Bacterial expression remains the most cheap and easy medium to scale up for protein production. At times the total protein produced by the cells, aggregates as inclusion bodies, where the target protein is incorrectly folded and non functional, this also makes it easier to be purified. For efficient recovery of correctly folded and functional recombinant proteins from inclusion bodies, optimization of protein refolding is a vital step. Several methods have been used such as high hydrostatic pressure [1], [2], [3], [4], use of chemical additives such as sugars, amino acids, polyols, alcohols etc [5] [6] [7], use of microfluidics chips [8],[9],[10]. Recent advances focuses on reducing protein aggregation during refolding, appraoches include on-cloumn refolding using artificial chaperones [10], [11], supression of aggregation with ion-exchange resins and small molelcule foldase mimics. The methods consider various parameters to be tweaked including protein concentration, redox conditions, temperature pH and ionic strength in order to optimize the refolding process. The principal idea behind protein refolding is gradual removal of denatured and addition of those compounds which enhances refolding and/or provide stabilization to the folded state. Traditionally dilution and dialysis have been used. Although every protein structure is unique to its properties, hence the optimum conditions vary with each protein and it has to be studied across array of refolding options and optimized accordingly.

Here we show protein refolding optimization of recombinant leucyl aminopeptidase (rPepL) of lactocaseibacillus casei expressed in inclusion bodies. The full-length rPepL polypeptide contains 411 amino acids including C-terminal 6X histag. In functionally active state it remains a homodimer, the monomer is of 42 kDa. The native structure catalyzes leu-*p*NA as substrate releasing a yellow-colored product, that has been used to monitor the efficiency of the buffer in refold denatured protein to its functionally active state. Molecular dynamics (MD) simulations in a mixed solvent system containing L-arginine, glycerol, and sucrose complemented these results revealing enhanced protein compactness and stabilization from the additives. The MD simulations provided atomistic insights into the effects of the mixed solvent environment on protein compactness, hydrogen bonding, radial distribution, and conformational energy landscapes.

## 2. Materials and methods

### 2.1 Protein expression

For protein expression 5ml of an E.coli strain (Lemo) overnight culture was used to inoculate 1L Luria-bertani broth. The cells were grown at 37 °C with shaking (225 rpm), 100µM of isopropyl-β-D-1-thiogalactopyranoside (IPTG) was added when the OD reached 0.4. and again, incubated for 5 hours. After that all the cells were harvested by centrifugation at 10000 g for 20 minutes at 10 °C.

### 2.2 Isolation of inclusion bodies

The cells were resuspended in 20mM Tris-HCl, and were sonicated at an amplitude of 53 kHz for 45 minutes. Cell debris along with soluble proteins were removed by centrifugation at 16000g for 45 min, resuspension of the pellet and sonication was repeated 3 times. Pellets containing inclusion bodies were washed with 20 ml of 20mM Tris-HCl, 500mM NaCl, 2M Urea, 2% Triton X100 pH 8.0, followed by centrifugation at 16000 g for 45 min, this step was repeated twice. Pellet were solubilized in 30 ml of 50mM Tris-HCl, 500mM NaCl, 8M Urea (pH-8.0) by overnight stirring at room temperature. Insoluble material were removed by centrifugation at 16000g.

### 2.3 Purification of PepL using denaturing conditions

For purification 2ml of the solubilized inclusion bodies of PepL was diluted in 10ml of binding buffer containing 50mM Tris-HCl, 500mM NaCl, 8M Urea, 20mM Imidazole (pH-8.0). This solution was applied to 2ml of Nickel-NTA resin column that was pre-equilibrated with the binding buffer. The protein solution was allowed to bind by shaking the column for 1 hour at room temperature. The bound protein was eluted using step gradient elution of Imidazole upto 500mM in the elution buffer. The purity of the elutes were analyzed with SDS-PAGE (12%). Pure elutes were pooled and concentrated using 10 kDa centricon (merck). The concentration of protein was determined by dividing the absorbance at 280nm in UV-2600 spectrophotometer (Shimadzu) by its extinction coefficient (67380).

### 2.4 Renaturation

The eluted denatured protein was refolded by drop wise dilution in various buffers. Protein concentration in each candidate buffer system was set to 0.6µM in 400ul of refolding buffer. The final concentration of urea from protein stock slotutions was set to 0.05 M. Refolding experiments were conducted in 12 well plates (costar; 150628), after adding the protein in the pre-cooled buffers, they were kept for for 3 hours at 4 °C on an orbital shaker. The samples were centrifuged at 16000 xg at 4 C to settle down aggregates and misfolds, the supernatant was used for activity and fluorescence experiments.

### 2.5 Chromogenic Leu-*p*NA Activity assay

The activity upon refolding were studied using chromogenic substrate Leu-*p*NA absorption coefficient of 9.9 M^-1^ cm^-1^ at 405nm. The reaction mixture contained 50mM phosphate buffer(pH-7.0), 2.5mM Leu-*p*NA, 1.0 mM CoCl_2_ and 0.1mg/ml of protein concentration in a final volume of 250 µl. The reactions were carried out on 96 well plates (Tarsons, India) at 25℃ for 1 hour. Absorbance were measured at 405 nm using Nanoquant plate reader (TECAN, Finland). For each assay two blanks were taken for the absence of protein and substrate in the reaction. The sum of the blanks were subtracted from the corresponding average of the test absorbance.

### 2.6 Fluorescence Spectroscopy probed with 1,8-ANS

A Stock solution of ANS was prepared with deionized water and its concentration was determined using the molar extinction coefficient of ε_350nm_ = 5000 M^-1^ cm^-1^. The solution of 2µM of protein was equilibrated with various molar ratios (20µM, 50µM, 200µM and 400µM) of ANS in 200µl of total solution. Fluorescence was recorded using a Cary Eclipse spectrofluorometer, Agilent Technologies. The measurements were performed using an excitation wavelength of 350 nm.

### 2.7 Mixed solvent Molecular dynamics

The dynamics and interaction of PepL 3D structure [12] in the optimum solvent composition of L-arginine, Sucrose and glycerol was determined using mixed solvent MD simulations. The simulation assesses structural stability, flexibility, solvent interactions, and energy landscapes through various computational analyses including RMSD, RMSF, RDF, PCA, and FEL. MD simulations of PepL in 0.5M L-arginine, 15% Glycerol and 10 % Sucrose was carried out for 100 ns in a cubic box of 200Å × 200Å × 200Å The final concentrations of the mixture compromised of 2409 molecules of Arginine, 9888 molecules of Glycerol and 1407 molecules of Sucrose. The mixture was solvated with 163755 molecules of water (TIP3P) at 350K. The GROMACS utility ‘insert-molecules’ was employed to introduce specified solute molecules: arginine, glycerol, and sucrose into the periodic boundary condition (PBC) simulation box. The number of molecules inserted for each solute was controlled using the *-nmol* option, allowing precise adjustment of solute concentration within the simulation environment. The parameter and topology files for the molecules were generated using the CGenFF (CHARMM General Force Field) server.

## 3. Results

### 3.1 Optimization of Refolding of purified and denatured inclusion bodies

The clean elutes were pooled and concentrated using 10 and 30 kDa ultrafiltration devices. The concentration for each protein was set upto almost 2.0 mg/ml and used as stock for the refolding experiments. 0.1 mg/ml was fixed for all refolding variations. All the refolding efficiency were compared to refolding efficiency Tris-HCl, as a control.

#### 3.1.1 Effect of L-arginine-HCl on refolding of Leucyl aminopeptidase (PepL)

Arginine has a charged guanidinium group (—C(NH2)(NH)NH2+) in the sidechain. Under biological conditions it assumes a net +1 charge. Arginine hydrochloride (ArgHCl) is created by reacting arginine with hydrochloric acid (HCl), producing a salt.

In the context of protein refolding studies, L-arginine-HCl has been extensively utilized [12] [13]. Optimal refolding of PepL was achieved at a 0.5 M concentration of L-arginine with refolding efficiency of 50.8%, with higher concentrations leading to reduced activity with refolding yield of 9%. At 1.5 M, activity dropped below levels observed without additives, indicating arginine’s role as a mild protein destabilizer Fig.1A [12]. Arginine stabilizes proteins against aggregation by slowing down protein-protein association reactions, without significantly affecting protein folding [14].

**Figure 1:**
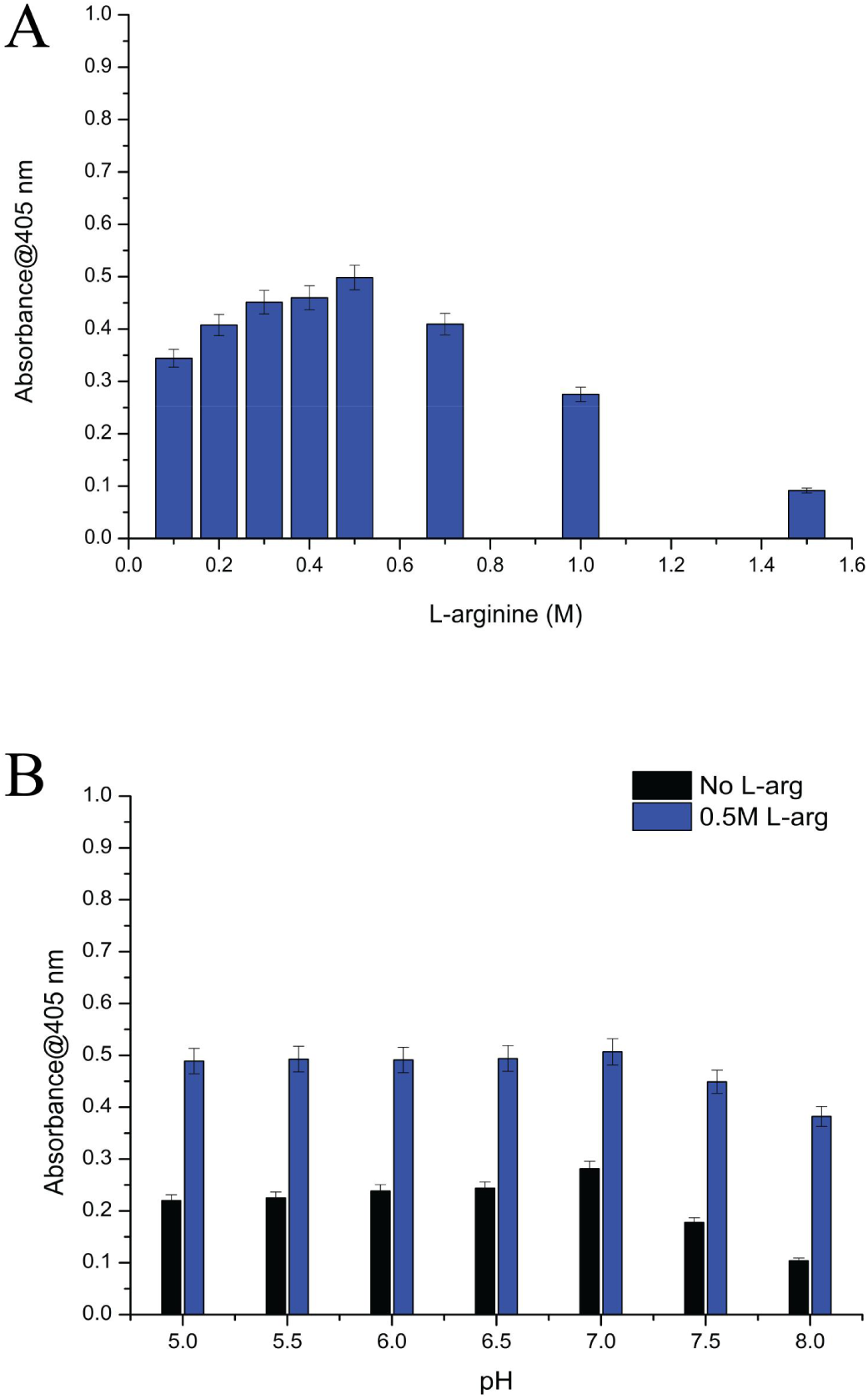
Refolding of 0.1mg/ml rPepL in various screens, monitored by Chromogenic Leu-pNA Activity assay. Refolding was conducted by dropwise dilution method at 4℃. (A) Various concentrations of L-arginine without any secondary additives. (B) Primary pH screen.

Maintaining a concentration of 0.5 M L-arginine we observed that increasing pH levels decreased PepL activity at pH 5 the refolding yield was 48% which increased to 50.6% at pH 7, and with a slower rate decreased to 38.2% at pH 8 (Fig:1B). Additionally, L-arginine showed additive effects with various sugars, polyols and osmolytes, although denaturants like GdmCl antagonized effects of Arg-HCl. Ishibashi and coworkers, 2005 demonstrated that L-arginine interacts differently with proteins compared to GdmCl. Lower concentrations of L-arginine exhibited negative preferential binding contrasting with GdmCl extensive protein binding behavior [15].

In this study, 0.5 M L-arginine alone demonstrated the highest efficiency in refolding compared to other additives studied individually; followed by 5% sarcosine, 5% glycerol, and then 15% sucrose. L-arginine exhibited additive effect when combined with other additives Fig.2

**Figure 2:**
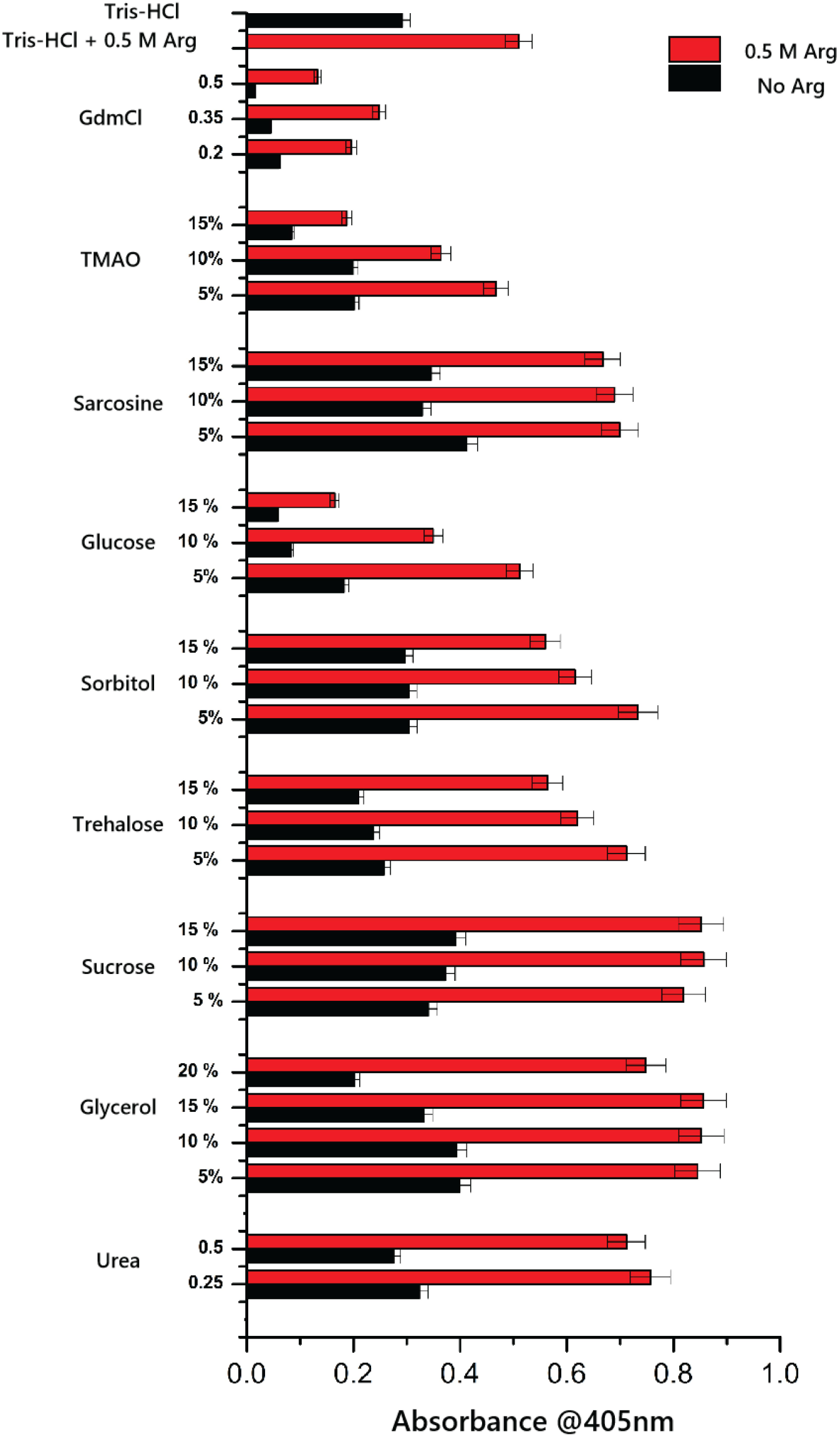
Composition of secondary additive screen. GdmCl,Urea and L-arginine are used in molar concentrations, rest of the additives were used in w/v ratio. Red Bars depict buffers with 0.5M L-arginine and Back bars depicts buffers without 0.5M L-arginine

#### 3.1.2 Effects of denaturants Urea and Guanidium hydrochloride

Removal of denaturants and assistance by co-solute or additives can prevent the unfolded proteins from getting aggregated and assists correct refolding, which is functionally active. We here observed that Urea in low concentrations promotes better refolding than GdmCl at 0.25 and 0.5 M Urea with a refolding yield of 32.3% and 27.4%, probably because urea might prevent aggregation more effectively during refolding [16]. GdmCl decreased refolding in concentration dependant manner from 5.9 % to 1.6% at 0.2 M to 0.5 M of GdmCl. Interestingly, in the presence of L-arginine, the yield increased to 19% at 0.25 M and 13% at 0.5M of GdmCl, but it lowered the efficiency of 0.5 L-arginine (Fig.2). This could be because GdmCl and Arg possessing highly denaturing guanidinium groups might be having additive denaturing affect because of total high concentration in combination. On the contrary urea showed increased refolding yield when added with 0.5M Arginine. At 0.25M and 0.5M Urea the refolding yields were 75.6% and 71.2 %, which is higher than control and 0.5M Arginine individually. Kohyama et al., 2010 showed refolding efficiency of 3SS lysozyme in which they decreased the urea and GdmCl concentration and increased the concentration of stabilizers [17].These show that urea at lower concentrations can assist protein refolding.

#### 3.1.3 Effects of Sugars on refolding of PepL

In this study the comparative effects of various sugars like Glucose, Sucrose, Trehalose was studied for enhancing the refolding yield of PepL. To observe the effects of Glucose, sucrose and trehalose three different concentrations were used 5% (w/v), 10% (w/v) and 15% (w/v). We observed that sucrose increased refolding yields from 34% (0.5M sucrose) to 39% (15%) with increase in concentration (Fig.2). With 0.5M L-arginine, sucrose increased the yield from 82% to 85% and the trend of refolding efficiency remained similar as observed in the absence of arginine. Sucrose showed additive effects with arginine and exhibited the maximal effect on refolding of PepL. Refolding yields in the presence of trehalose and glucose were lower than sucrose. The refolding yield decreased with increasing concentrations of trehalose and glucose. Trehalose enhanced refolding by 25.6% at 5% to 20.9% at 15%. Similarly, glucose with lower yield than trehalose increased refolding by 18% at 5% to 5.6% at 15% (Fig:2). Although trehalose is well known for its better stabilizing effect on protein [18]. The lower refolding yield of PepL could be owing to formation of PepL aggregates at 10% and 15 % of trehalose (without arginine). Although trehalose showed additive effect when used in combination with 0.5M L-arginine. Upon addition of arginine the yield increased up to 71.18% at 5%, but it steeply declined to 56% at 15% trehalose. This suggests that although trehalose improved refolding of PepL, yet it induced PepL aggregation at higher concentration, which could not be completely off-set by L-arginine. Glucose has negative effects on refolding of PepL as it lowers the refolding yield than Tris-HCl.

#### 3.1.4 Effects of osmolytes on PepL refolding

In this study we used two osmolytes Tri-methylamine N oxide (TMAO) and sarcosine. TMAO is an organic compound, neutral, while sarcosine is a amino acid derivative, neutral. Both the compounds are known to increase stabilization of proteins [19] [20]. Although TMAO reduces refolding yield than control, at three concentrations 5%, 10% and 15% the refolding yield went down from 25.6%, to 20.9% with increasing TMAO concentrations (Fig.2). Yi-ting liao and coworkers demonstrated that TMAO reduces the surface tension of water but stabilizes protein by acting as a surfactant [19]. TMAO also reduced the positive effects of L-arginine as the refolding yield was 46.6% which is maximal at 5% TMAO + 0.5M L-Arg and 36.4 % and 18.7 % at 10% and 15 % of TMAO with 0.5M L-Arg. The TMAO concentration dependence decrease in refolding yield of PepL suggested that TMAO inhibit the refolding of PepL.

Although sarcosine has not been widely used in protein refolding as an additive, but it has been used to stabilize lysozyme [21]. We used sarcosine at 5% 10% and 15% concentrations, with highest refolding yield of 41.2% at 5% which further reduced to 32 % at 10% and 34 % at 15% of sarcosine. Lowest concentrations of sarcosine promote refolding better than its higher concentrations. With 0.5M L-Arg, 5% sarcosine showed highest refolding yield of 69.9% that decreased to 66% at 15%. It shows an additive effect of sarcosine on PepL refolding when used in combination with 0.5M L-Arg.

#### 3.1.5 Effect of Polyols

In this study we used two polyols, namely glycerol and sorbitol. Glycerol is a 3C polyol, and sorbitol is a 6C polyhydric alcohol. Both increased the refolding yield of PepL. Glycerol is known to enhance hydrophobic interactions by increasing solvent ordering around the protein molecule and also by preventing aggregation during refolding [22] [23]. Sorbitol is a stabilizer of proteins by increasing preferential hydration towards hydrophilic regions of the protein [24]. Glycerol showed higher refolding yields than the control. Glycerol promoted the highest refolding yield of 39.9 % at 5%, but its increasing concentration reduced refolding yield. At 20% glycerol the refolding reduced to 20%, which is lower than the control. But it showed much higher yield in the presence of 0.5M L-arg. The refolding yield doubled upto 84.5% at 5% glycerol, and 85.19 % and 85.5% at 10% and 15 % of glycerol (Fig.2). A concentration of 5-15% glycerol showed almost similar effect on refolding yield in the presence of L-arginine. However, the yield was 74.8% at 20 % glycerol. In comparison, sorbitol showed almost no effect on refolding yield at 5-15% in comparison to control (Tris-HCl) however, when combined with L-arginine, the overall yield increased at 5-10% and then no effect at 15% in comparison to control (0.5M L-arg). Sorbitol also showed a concentration dependent decrease in refolding from the maximum achieved at 5% sorbitol, suggesting lower concentrations assist rPepL refolding.

It seems that rather than total content of carbons and-OH groups, the molecular structure of solute molecular is important in imparting effect on refolding. Glycerol has 3 carbon and 3 hydroxyl groups and in term of %age concentration, while sorbitol has 6 carbon and 6 hydroxyl groups. Thus, they contribute almost equally in term of chemical group at a given concentration in %age in a solution. However, their effect is not similar, suggesting the molecular structure of the solute is also important in imparting their effect on properties of bulk, eg, glycerol reduces the surface tension of water [25]while sorbitol increases [26] This should effect the hydration of solute itself and of protein that should in turn cause different salting out effect and of proteins refolding at higher concentration. It has been also observed that some co-solvents like glycerol and other higher polyols (4-6 carbons) behave in a protein-specific manner in refolding and stability of protein.

### 3.2 1,8-ANS probed Fluorescence Spectroscopy

An equilibrated solution of 2µM protein with 100X ANS were prepared and the fluorescence spectra was recorded. (Fig.3) shows that with 0.5 M arg+ 10% sucrose the fluorescence spectra was highest among the denatured (8M urea) and 0.5 M urea. There is a blue shift in the peak of the spectra when concentration urea is reduced from 8M to 0.5M, the shift depicts the helical increment upon refolding. A majority of the population of the protein molecules have been refolded to their native state in 0.5 M arg+ 10% sucrose, corresponding with its higher activity.

**Figure 3:**
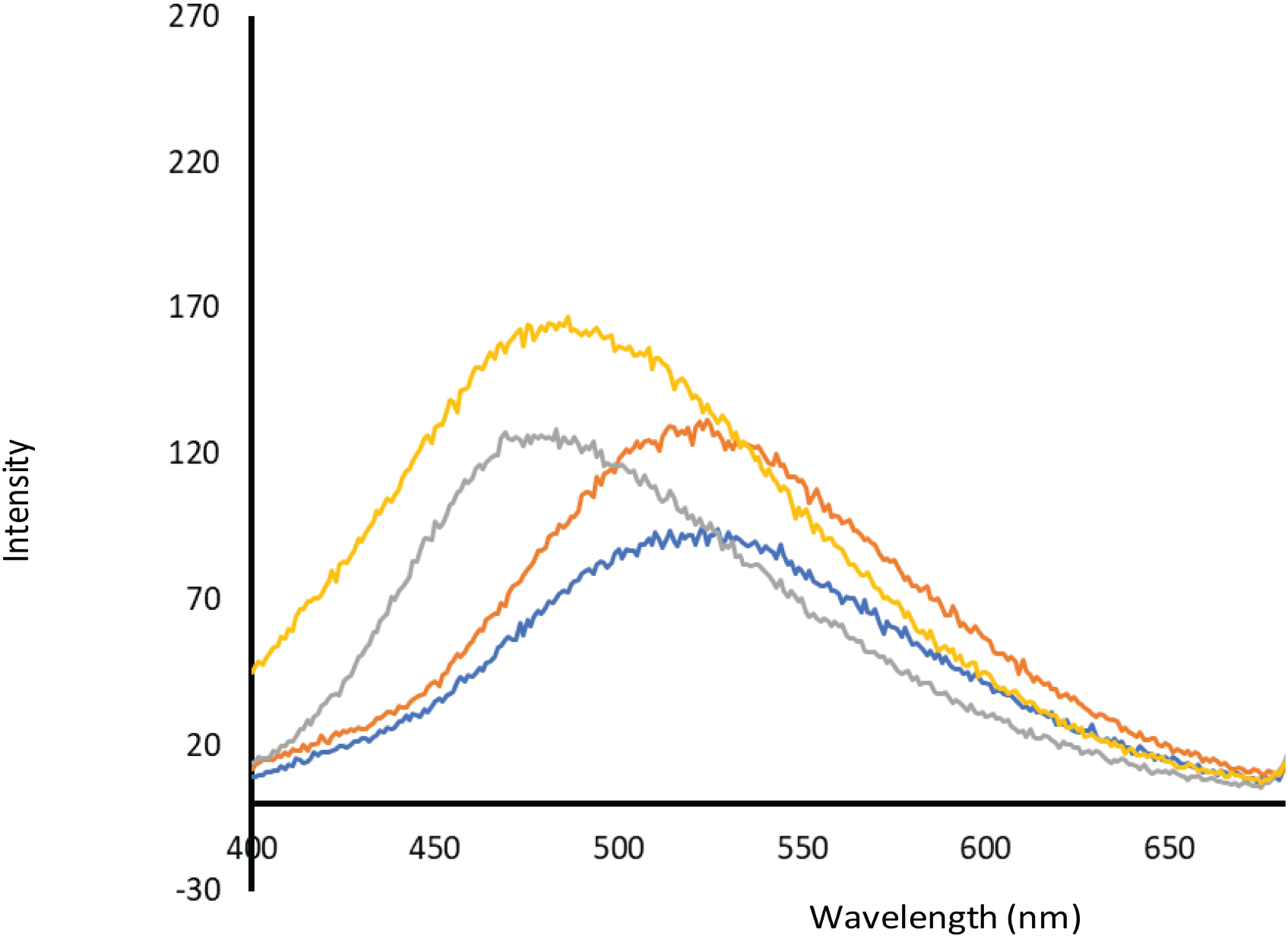
Fluorescence Spectroscopy probed with 1,8-ANS. Concentration for 1,8-ANS to attain best resolution was optimized. 0.1 mg/ml rPepL were mixed with 2uM 100X 1,8-ANS on ice and kept for equilibration before reading. Blue Line depicts blank, Red depicts 8M Urea, Grey depicts 0.5M L-arginine (without any secondary additives) and Yellow depicts 0.5 M L-arginine + 15% sucrose + 10% Glycerol respectively. Blue shift in the spectra can be seen the optimized refolding buffer.

### 3.3 Mixed Solvent Simulations

We have performed MD simulations of PepL in a equilibrated solution of L-arginine, glycerol and sucrose at a fixed concentration for 100ns. In order to understand nature of interactions through which the solvent stabilizes the native state making it less prone to aggregation in the experimental conditions, the root mean square deviation (RMSD), number of hydrogen bonds, radial distribution function (g(r)), Principal component analysis (PCA) and free energy landscape (FEL) were calculated.

The structural conformation in mixed solvent was stable and exhibits compactness in the dynamics of PepL relative to simulations in water. The mixed solvent trajectory readily achieves an equilibrated conformation which doesnot deviate much throughout 100ns, and exhibits an average RMSD of 0.35 nm, while in water PepL attained an equilibrated conformation at around 40ns, with average RMSD of 0.47 nm Fig.4.

**Figure 4:**
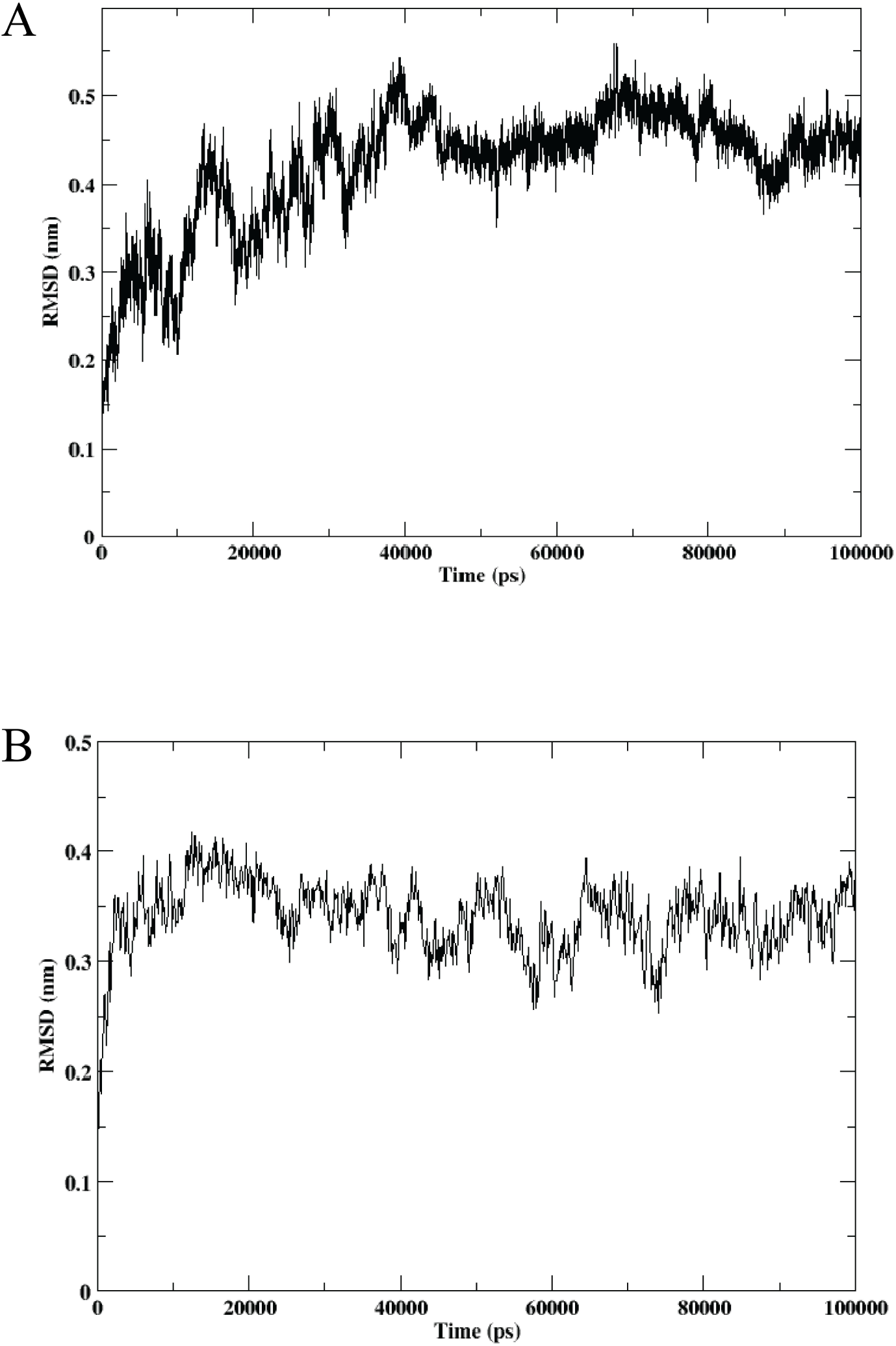
Cα-RMSD of PepL over 100 ns at 350K in (A) Water (B) Mixed solvent solution of L-arginine, Sucrose and Glycerol.

#### 3.3.1 Hydrogen bonds

The hydrogen bond formed over a 100ns between the protein and co-solvents (Arginine, Glycerol, and Sucrose) are given in the Fig.5. Among the three, the hydrogen bond between protein-glycerol consistently forms the highest number of hydrogen bonds, fluctuating mostly between 50 and 85, indicating strong and frequent interactions with the protein. Further protein-sucrose maintains a moderate number of hydrogen bonds, generally between 20 and 35, suggesting relatively stable but less intense interaction. In contrast, the hydrogen bond between protein-arginine forms the fewest bonds, fluctuating between 5 and 20, indicating weaker or less frequent interactions. Overall, glycerol interacts more in comparison to sucrose and arginine exhibits the strongest hydrogen bonding with the protein.

**Figure 5:**
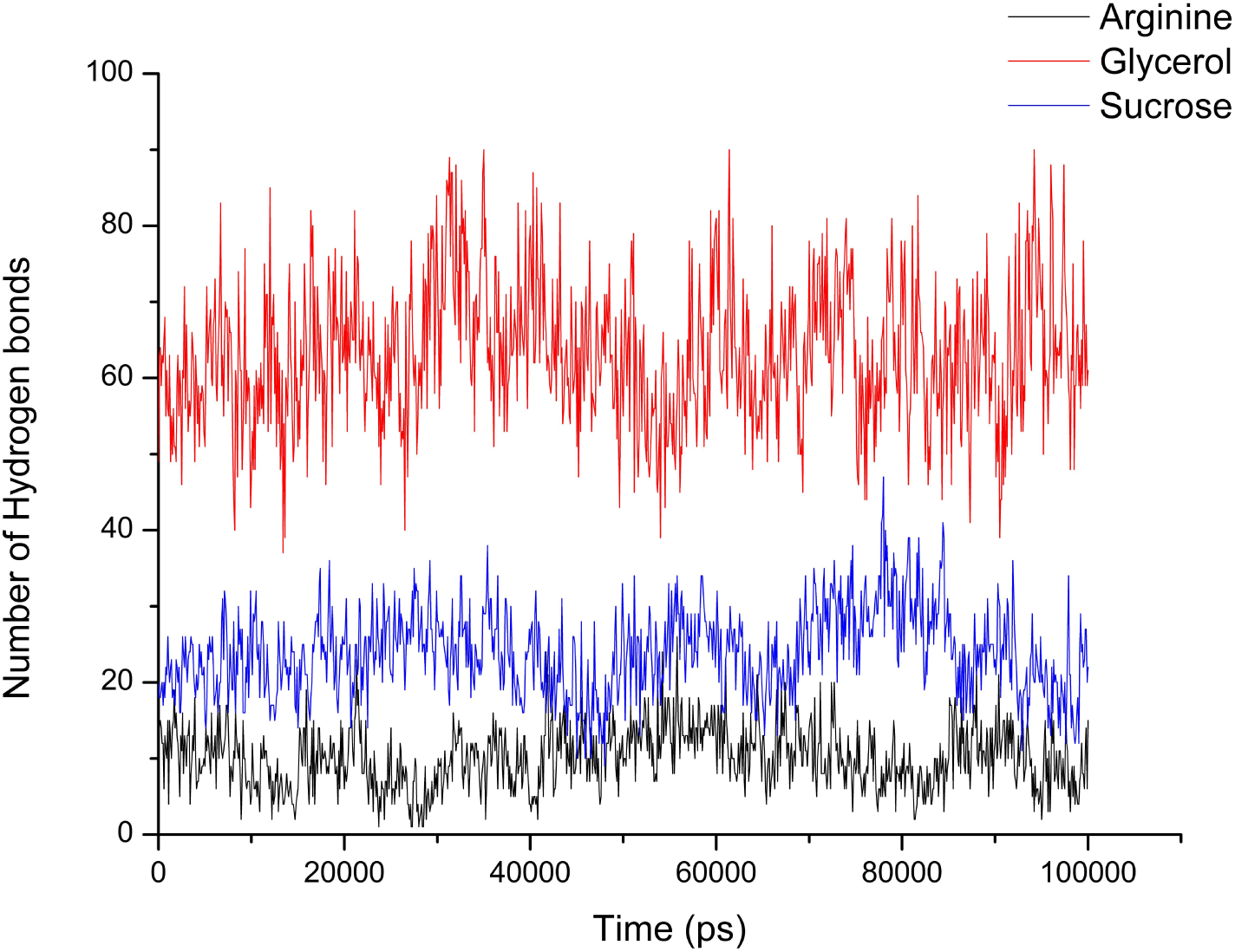
Hydrogen bond formed between protein and the molecules during the 100 ns of simulation

#### 3.3.2 Radial Distribution Function

To elucidate the molecular interactions and spatial organization of the cosolvents around the protein, radial distribution function (RDF, g(r)) analyses were performed. The first RDF plot (Fig.6) represents the cosolvent–cosolvent distribution, in the simulated systems.

**Figure 6:**
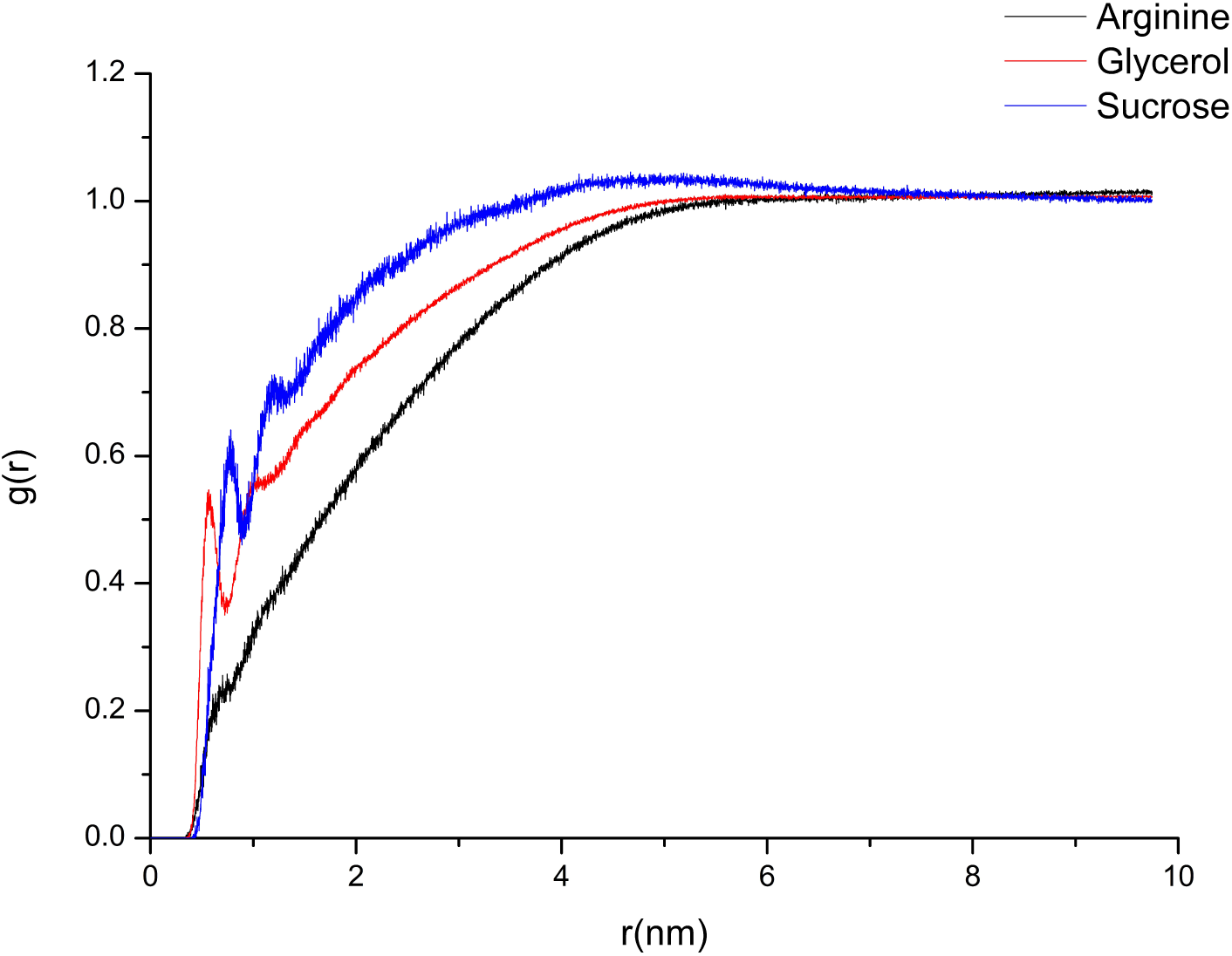
Radial distribution function (RDF) plot showing the spatial distribution of glycerol, arginine, and sucrose molecules around the protein, using the center of mass of each molecule as the reference point.

In the cosolvent–cosolvent RDF profile (Fig.6), sucrose exhibited a pronounced first peak around 0.5–0.8 nm, indicating a strong local ordering and extensive hydrogen-bonding network among sucrose molecules. Glycerol showed a moderate and broader first peak, reflecting intermediate self-association, whereas arginine displayed a relatively flat and diffuse distribution, suggesting a more homogeneous dispersion in the solvent phase. The higher structuring in sucrose implies strong intermolecular cohesion and limited mobility, consistent with its known ability to form a compact hydration matrix around biomolecules. Although glycerol molecules exhibited closer intermolecular association in the cosolvent–cosolvent RDF, their lower g(r) near the protein surface indicates reduced population around the protein, consistent with preferential hydration rather than direct surface binding.

#### 3.3.4 Principal Component analysis

The conformational variations are restricted to 4 nm (PC1) in the case of mixed solvents and spans much wider in water 20nm in (PC1) Fig.7 A,B. The data points form two distinct clusters in mixed solvents, indicating that the protein samples at least two major conformational states or energy minima during the simulation. The separation along the eigenvector 1 axis suggests significant structural transitions between these states. The density and spread of the points within each cluster reflect the extent of conformational flexibility within each state. Conformational displacements are higher in water as compared while in mixed solvents we see moderate values. Overall, this PCA analysis suggests the presence of conformational heterogeneity and transitions between stable states during the simulation.

**Figure 7:**
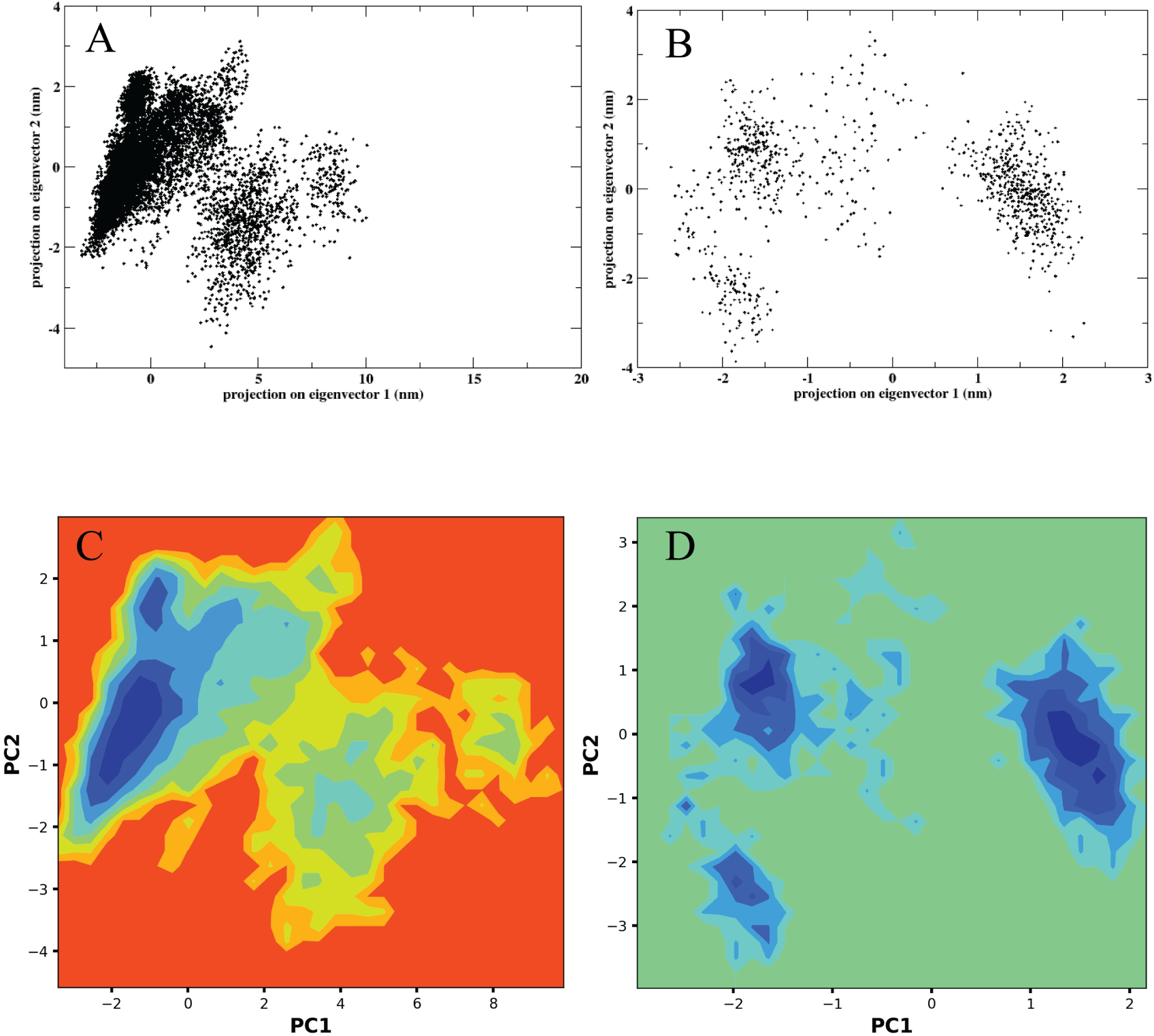
PCA representing the conformational space sampled by the protein during the molecular dynamics simulation (AB). 2D representation of Free Energy Landscape throughout 100 ns simulation (CD). (A, C) Water, and (B, D) Mixed solvent.

#### 3.3.3 Free Energy landscape

The Free Energy Landscape (FEL) plot shown in the Fig.7 C,D derived from Principal Component Analysis (PCA), illustrates the conformational space sampled by the protein during the molecular dynamics simulation, with PC1 and PC2 representing the dominant motions. The color gradient from dark blue to green indicates increasing free energy, where dark blue regions correspond to low-energy, thermodynamically stable conformations, and green areas represent higher-energy, less stable states. The presence of multiple distinct low-energy basins suggests that the protein adopts several stable conformations throughout the simulation, with transitions between them requiring energy to overcome intermediate barriers. This landscape highlights the dynamic nature of the protein and provides insight into its structural flexibility and potential functional motions.

The finding is similar as the FEL exhibits similar span as PCA plots. PC1 caputres much varied conformations in water. Here it shows that 3 conformations which have the least free energy the plot shows three basins eventually population low energy conformers on contrary in water, PepL attains a single basin and explores much larger conformational space in the energy landscape. These results provide insights into the differential influence of small solute molecules on the structural dynamics and stability of the protein under simulated physiological conditions.

## 3. Discussion

### 4.1 Experimental refolding of rPepL is enhanced with the use of L-arginine in combination with secondary additives

Expression of rPepL in reasonable quantities requires large scale production in E.coli. which mostly generates inclusion bodies (Ibs). These Ibs later could only be purified in denaturing conditions here it is in 8M Urea. To recover substantial amount of soluble active protein a strategy was needed to efficiently refold the purified but denatured rPepL. This being the first attempt to evaluate refolding of rPepL with various chemical additives, we used chemicals of varied nature such as, sugars, osmolytes, polyols, amino acid and denaturants. The additives were used both alone and in combination at a fixed pH, ionic strength, temperature and residual denaturant i.e. 0.1M Urea. We conducted low temperature (4 ℃) drop-wise dilution method for all the refolding experiments. We found that 0.5 M L-arginine-HCl was the most effective additive for refolding Leucyl aminopeptidase (PepL), achieving a refolding yield of 50.8%, with higher concentrations being destabilizing Fig 1 A. Optimal refolding occurred at pH 7 Fig. 1 B, and L-arginine displayed additive effects when combined with stabilizers like sucrose, glycerol, and sarcosine, significantly boosting yields (up to 85% with sucrose) Fig.2. Urea enhanced refolding more than GdmCl, especially when paired with arginine, while GdmCl antagonized L-arginine’s effects. Trehalose and glucose had less pronounced benefits, with glucose showing a negative impact. TMAO reduced refolding efficiency, and glycerol showed strong performance, particularly in combination with arginine, while sorbitol had minimal impact on its own. The optimal combination for refolding rPepL is 0.5 M L-arginine-HCl combined with 5-15% sucrose and 5-15% glycerol Fig.2. This mixture leverages L-arginine’s stabilizing effects and enhances refolding efficiency, achieving yields up to 85%. Sucrose and glycerol complement L-arginine by improving protein stability and preventing aggregation, making this combination the most effective for maximizing refolding yields.

The use of L-arginine-HCl in protein refolding studies has been widely explored due to its unique properties in stabilizing proteins against aggregation and facilitating proper folding. The optimum concentration of 0.5M balances the stabilizing effects of arginine without inducing protein destabilization, as observed with higher concentrations where refolding yields decreased significantly. This phenomenon has been attributed to arginine’s ability to modulate protein-protein interactions, thereby preventing non-native aggregation pathways while maintaining the native folding pathway [13]. Previous studies have reported that arginine efficiently refolds proteins by various mechanism such as, its guanidinium group interacts with tryptophan side chains of the protein that suppress aggregation [27], it binds to the denatured protein mainly through hydrogen bonding excluding itself gradually as the protein recovers its native conformation and it protects some amide deuterium atoms from being exchanged with hydrogen during protein refolding process within the proteins α-helix domain which reversibly stabilizes denatured and partially folded state [28] and it weakens protein-protein interaction rather than stabilizing the protein, stated as the main reason for arginine suppressing protein aggregation [29]. Moreover, our results corroborate with findings that L-arginine interacts differently with proteins compared to denaturants like guanidinium chloride (GdmCl). L-arginine exhibits a mild destabilizing effect on proteins, primarily affecting protein-protein association reactions rather than the folding process itself [15]. This is in contrast with the effect of GdmCl, which extensively binds to proteins and unfolds them at higher concentrations. The interaction between GdmCl and L-arginine, as observed in our study, suggests a relationship where the denaturing effects of GdmCl might be mitigated to some extent by L-arginine. While GdmCl typically decreases refolding efficiency, the presence of L-arginine led to increased yields (19% at 0.25 M and 13% at 0.5 M GdmCl) Fig.2, albeit with reduced efficiency at higher concentrations of L-arginine (0.5 M). This could imply that the guanidinium groups in both GdmCl and L-arginine may have additive denaturing effects when their concentrations are high.. The additive effect observed with urea and L-arginine further supports the notion that these additives can complement each other in promoting protein refolding. At 0.25 M and 0.5 M urea concentrations, refolding yields reached 75.6% and 71.2%, respectively, surpassing those achieved with either urea or L-arginine alone. This effect is consistent with findings by Kohyama et al. (2010), who demonstrated enhanced refolding efficiency of 3SS lysozyme by decreasing urea and GdmCl concentrations while increasing stabilizer concentrations [17].

Sucrose plays a crucial role in protein refolding and stabilization by acting as a chemical chaperone, preventing aggregation and aiding in the recovery of denatured proteins. It stabilizes the native structure of proteins against chemical denaturants and heat by increasing packing density and serving as a crowding agent. In environments with other crowding agents like dextran, sucrose helps counterbalance their aggregation-promoting effects, leading to improved refolding rates and yields. Additionally, sucrose promotes the compaction of both folded and unfolded protein states, reducing conformational entropy and enhancing overall protein stability [30] [31] [32]. Our observation of sucrose with and without L-arginine in enhancing PepL refolding efficiency is consistent with these reports. Without arginine, sucrose has maximum refolding efficiency of 39%, that is because of its stabilizing effect, although it might not have suppressed aggregation like arginine, but when arginine is supplemented which have lowered aggregation, sucrose showed an additional rise in the refolding efficiency upto 85% benefiting from the dual effect of arginine and sucrose. Glycerol has also been reported for high refolding efficiency and stabilization of protein [33], [34], [35]. It facilitated lysozyme refolding by stabilizing partially folded intermediates and reduced the activation barrier for unfolding, Chen and coworkers showed that the nature of protein determines the effect of glycerol, with hydrophobic-dominated proteins showed increased stability [36]. We see that with higher concentration of glycerol the refolding decreases, this is similar to with the findings by W.Ou and coworkers [37], refolding yield of creatine kinase was better at 25% glycerol, but with high concentration the yield reduced. Preferential hydration, suppressing aggregate formation, and increasing secondary structure propensity can be the reason for its relatively better refolding efficiency. When added with arginine all the positive attributes from both the compounds come into a synergistic play and increase refolding efficiency upto 85.5%, but then again at higher concentration of glycerol (at 20%) the yield decreases, suggesting a balance between the abundance of the molecules, overcrowding may incur decrease in refolding efficiency. When only glycerol was used from 5% to 20% the refolding yield decreased, but with arginine the decline is only seen at 20%, shows that arginine was compensating for the negative effects of high concentration of glycerol upto 15%. Although Trehalose is recognized for its ability to stabilize protein, by maintaining protein structure and preventing aggregation at lower concentrations [18] conversely here its refolding efficiency is lower than that of Tris-Hcl only. Without arginine, we observed aggregation in all the used trehalose concentrations, with increasing concentrations of trehalose arginine could not compensate the effects over increasing concentration is because of their union, we see decrease in yield with increasing trehalose concentration when used with arginine. Similarly glucose had much lower yields and showed decline at higher concentrations, same. Glucose binding can stabilize protein structure and slow down unfolding-refolding process [38]. Trimethylamine N oxide (TMAO), despite its known stabilizing effects on proteins in their native states, showed a concentration-dependent decrease in refolding yield of PepL in this study. This observation is consistent with findings by Yi-ting Liao and colleagues, who highlighted TMAO’s dual role in reducing the surface tension of water and stabilizing proteins by acting as a surfactant [19]. However, our results indicate that TMAO inhibits the refolding process of PepL, as evidenced by decreased refolding yields at higher concentrations (20.9% at 10% and 15% TMAO). Furthermore, TMAO mitigated the positive effects of L-arginine, reducing refolding yields significantly when combined with L-arginine at higher TMAO concentrations (36.4% and 18.7% at 10% and 15% TMAO with 0.5M L-Arg). This suggests a potential interference or competition between TMAO and L-arginine in their mechanisms of protein stabilization during refolding. In contrast, sarcosine, an amino acid derivative less commonly used in protein refolding studies, demonstrated interesting effects. While higher concentrations of sarcosine (10% and 15%) reduced PepL refolding yields, lower concentrations (5%) showed the highest refolding yield (41.2%). This aligns with previous studies where sarcosine has been shown to stabilize proteins like lysozyme under certain conditions [21]. Moreover, when combined with 0.5M L-arginine, 5% sarcosine exhibited an additive effect, enhancing PepL refolding yield to 69.9%. This additive effect suggests that sarcosine may complement the stabilizing effects of L-arginine, potentially by modulating protein-water interactions or surface properties. In contrast, sorbitol, a 6-carbon polyol, showed minimal effect on rPepL refolding yield when used alone at 5% to 15% concentrations, compared to the Tris-HCl control. However, when combined with L-arginine, sorbitol demonstrated an increase in refolding yield at lower concentrations (5% to 10%), suggesting a supportive role in protein stabilization similar to glycerol. Nevertheless, higher concentrations of sorbitol (15%) resulted in no significant improvement in refolding yield compared to L-arginine alone, indicating a concentration-dependent effect where sorbitol may not enhance refolding efficiency beyond certain thresholds.The blue shift observed in the fluorescence spectra upon reducing urea concentration from 8 M to 0.5 M is consistent with previous studies indicating the reorganization of protein structure towards its native conformation. This shift reflects the helical increment and decrease in exposed hydrophobic surfaces that bind ANS, confirming the structural changes corresponds to formation of secondary structure [39]. Spectra of samples in 0.5M arg + 10% sucrose have an incresed area under the curve implying more amount of protein has been refolded which is also in agreement with its activity Fig.3.

### 4.2 Molecular dynamics in mixed solvents

In this work we performed MD simulations to explore the efficiency of 0.5M L-arginine, 15% glycerol and 10% sucrose to stabilize the native folded form at 350K Mixed solvent combination exhibited stabilization to the structure, in the overall 100 ns trajectory the PepL was more compact and traversed less through varied conformations in comparison to water. Through simulations we get a hollistic ovreview of the mechanism through which compounds in the solution have interacted eventually providing stabilization to the structure at temperature than higher PepL melting temperature. Higher kinetic velocity will resolute the findings better than conducting it at lower temperatures. We find that for PepL stabilization two different mechanisms come into play (1) supression of aggregate formation and (2) stabilization of the native folded structure. 0.5M L-Arginine interacts less with the structure as its g(r) is the lesser than glycerol and sucrose, the shorter hydrogen bonds formed by arginine reflects its direct electrostatic and bifurcated interactions between the guanidium group and the negatively charged residues such as ASp and Glu, which are transient yet localized with the structure in comparison to other two additives. thereby suppressing non-native protein-protein associations that lead to aggregation. At higher concentrations, these interactions become excessive, leading to self-association of arginine molecules, disrupting solvent structure and reducing refolding efficiency—a behavior previously reported by [15],[16],[40]. reported that with higher contration of arginine the structuration of arginine molecules around the protein reduces as they tend to form clusters. Its lowest g(r) indicates minimal residence time compared to glycerol and sucrose. This implies that arginine’s role in terms of increasing refolding effiecncy arises not from persistant surface binding but from dynamic sheilding, as it interacts transiently to neutralize local charges and then disperses, stabilizing the proteins against aggregation [15].

Glycerol significantly improved refolding, especially when combined with arginine, where yields reached 85%. As a polyol, glycerol tends to stay away from the protein surface, which increases the chemical potential of water molecules near the protein and encourages the compact native shape.This effect, known as preferential hydration, boosts thermodynamic stability by favoring the folded state. Hydrogen bonding analysis from MD simulations showed that glycerol formed the most hydrogen bonds with the protein, ranging from 50 to 85 over 100 ns. This reflects its strong stabilizing interactions and its ability to structure the solvent. Furthermore, the consistent and stable RMSD profiles seen in mixed solvents suggest that glycerol helps keep the protein structure compact and balanced. These findings support previous experimental and computational studies that show glycerol can reduce aggregation and stabilize folded proteins by improving water structuring and strengthening the solvation layer [23],[35].

Sucrose, which exhibits preferential exclusion behavior and has a characteristic to prefer the excluded phase, contributed supplementary stabilization to rPepL. In experiments, sucrose elevated refolding efficiency in a concentration-dependent manner with a maximum at 10–15% (w/v), where it combined with arginine in a synergistic manner to obtain up to 85% refolding. RDF profile calculated from the results of MD simulation demonstrated that the highest spatial association (g(r)) of sucrose was localized close to the protein surface, which indicates the development of a clear hydration shell lowering the local water mobility and restricting solvent penetration inside the protein core and hence forestalls partial unfolding.

The function of sucrose as a thermodynamic stabilizer encompasses the enhancement of the free-energy disparity between the folded and unfolded states through its preferential exclusion from the protein surface. This process significantly diminishes the conformational entropy associated with the unfolded state, thereby favoring the equilibrium position towards the native conformation [31]. The moderate level of hydrogen bonding detected during the simulation (20–35 hydrogen bonds) provides additional evidence for a model in which sucrose facilitates surface protection without excessive interactions, which is consistent with the concept of “soft stabilization” observed in other refolding investigations [30].

Arginine (0.5 M), glycerol (15%), and sucrose (10%) in combination was superior to all single additives and also gave highest refolding yield of ∼85%. Their combined synergy comes from due to their collectively facilitating mechanisms:

1. Arginine inhibits aggregation by transient electrostatic interaction.
2. Glycerol enhances native-like hydrogen bonding networks and structural compaction.
3. Sucrose contributes to osmotic stabilization and hydration-shell reinforcement.

Simulations of the mixed solvent confirmed this synergy. The protein was stable in its RMSD (∼0.35 nm), consistent with a stable and compact conformation. Free energy landscape (FEL) had several basins of low energy, typical of conformational stability and access to energetically optimal native states. Principal component analysis (PCA) also confirmed limited conformational sampling relative to water-based simulations and substantiates the stabilizing influence of the mixed solvent environment. In summary, these results indicate that the ternary mixture accomplishes protein stability and conformational breathing required to obtain productive rather than kinetic trapping during protein refolding.

## 5. Conclusion

In conclusion, the optimization of refolding conditions for recombinant Leucyl aminopeptidase (rPepL) demonstrated that a combination of L-arginine, glycerol, and sucrose yielded the highest refolding efficiency among all the other additives. L-arginine played a crucial role in suppressing aggregation and enhancing protein stabilization, showing synergistic effects when combined with other additives such as glycerol and sucrose. The additives used in our experiments contributed to both aggregation suppression and protein stabilization, two critical factors for successful refolding. L-arginine is particularly effective at preventing aggregation, as it disrupts protein-protein interactions during the folding process. This is essential because aggregation often occurs when unfolded proteins misfold and clump together, reducing the yield of properly folded protein. L-arginine’s ability to shield charged surfaces also promotes refolding by reducing intermolecular interactions. Meanwhile, stabilizing additives like glycerol and sucrose contribute primarily by enhancing the structural stability of the folding protein. These molecules increase the preferential hydration around the protein, which enhances its native conformation by promoting proper hydrophobic packing. Glycerol and sucrose are known to increase the thermal stability of proteins by stabilizing the folded state, making it less prone to denaturation. The dual role of these additives is evident in their synergistic effects. L-arginine prevents aggregation, allowing proteins to fold correctly without interacting with other misfolded proteins, while glycerol and sucrose enhance the proper folding by stabilizing the protein’s native structure. This combined action improves refolding efficiency, resulting in a higher yield of functional proteins. This dual contribution is critical to achieving the balance necessary for both preventing misfolding-related aggregation and ensuring the protein reaches a stable, functional conformation.

Urea at lower concentrations also improved refolding, while higher concentrations of GdmCl inhibited the process. The observed blue shift in fluorescence data indicated that the protein achieved a more compact, folded conformation, correlating with improved refolding yields.

Molecular dynamics simulations provided atomistic insight into the synergistic stabilization mechanisms of these additives. Glycerol formed the largest number of hydrogen bonds with the protein, reinforcing hydration layers and compactness; sucrose contributed through preferential hydration and formation of an osmotic protective shell; and arginine transiently interacted through electrostatic and bifurcated hydrogen bonds, reducing aggregation-prone contacts. Radial distribution and coordination analyses demonstrated that each cosolvent stabilizes the protein via distinct yet complementary interactions, while PCA and FEL analyses confirmed restricted conformational fluctuations within stable, low-energy basins.These findings provide insights into the effective use of additives for enhancing protein refolding from denatured inclusion bodies.

## Contribution of authors

1. Conceptualization: DD and JKK
2. Data curation: DD
3. Investigation: DD
4. Methodology: DD
5. Analysis: DD
6. Project administration: JKK
7. Resources: DD and JKK
8. Writing-review and editing: DD and JKK
9. Funding Acquisition: DD and JKK

## Declaration

Authors declare no conflict of interest

